# Juvenile corals inherit mutations acquired during the parent’s lifespan

**DOI:** 10.1101/2020.10.19.345538

**Authors:** Kate L. Vasquez Kuntz, Sheila A. Kitchen, Trinity L. Conn, Samuel A. Vohsen, Andrea N. Chan, Mark J. A. Vermeij, Christopher Page, Kristen L. Marhaver, Iliana B. Baums

## Abstract

128 years ago, August Weismann proposed that the only source of inherited genetic variation in animals is the germline^1^. Julian Huxley reasoned that if this were true, it would falsify Jean-Baptiste Lamarck’s theory that acquired characteristics are heritable^2^. Since then, scientists have discovered that not all animals segregate germline cells from somatic cells permanently and early in development^3^. In fact, throughout their lives, Cnidaria^4–6^ and Porifera^7^ maintain primordial stem cells that continuously give rise to both germline and somatic cells. The fate of mutations generated in this primordial stem cell line during adulthood remains an open question. It was unknown whether post-embryonic mutations could be heritable in animals^8–10^—until now. Here we use two independent genetic marker analyses to show that post-embryonic mutations are inherited in the coral *Acropora palmata* (Cnidaria, Anthozoa). This discovery upends the long-held supposition that post-embryonic genetic mutations acquired over an animal’s lifetime in non-germline tissues are not heritable^2^. Over the centuries-long lifespan of a coral, the inheritance of post-embryonic mutations may not only change allele frequencies in the local larval pool but may also spread novel alleles across great distances via larval dispersal. Thus, corals may have the potential to adapt to changing environments via heritable somatic mutations^10^. This mechanism challenges our understanding of animal adaptation and prompts a deeper examination of both the process of germline determination in Cnidaria and the role of post-embryonic genetic mutations in adaptation and epigenetics of modular animals. Understanding the role of post-embryonic mutations in animal adaptation will be crucial as ecological change accelerates in the Anthropocene.

## Main

With the exception of planarian flatworms^11^, bilaterian animals segregate germline cells from somatic cells early in development^3^ (Fig. 1a). Because most animals segregate germlines early in development, it has long been assumed that only germline mutations are inherited in animals. Thus, genetic mutations that occur after early development (*i.e.*, post-embryonic mutations in the somatic tissues) cannot be inherited in these animals—limiting their evolutionary impact. In contrast, plants segregate germline cells late in development^12–14^ and pre-germline mutations are heritable. According to the genetic mosaicism hypothesis^15–17^, such post-embryonic mutations provide genetic diversity for adaptation to local conditions^16^. At the base of the metazoan tree, sessile cnidarians share life history characteristics with plants, including modular growth^18^, long lifespans^19^, high capacity for regeneration^20^, continuous germline determination^3^, and alternating asexual/sexual reproductive cycles^21,22^. One group of cnidarians, scleractinian corals, exemplify many of these characteristics. Scleractinian coral species often reproduce by asexual cloning and fragmentation and some species have remarkably long lifespans, estimated to be upwards of thousands of years^19^, allowing genets to reproduce for centuries or even millennia. However, projected environmental changes could lower the fitness of these previously well-adapted genets. Scleractinian corals and other sessile colonial animals may instead acquire mutations during adulthood^3^ and subsequently pass these mutations onto their offspring (Fig. 1); some of these mutations may be beneficial and contribute to adaptation. However, continuous germline determination has not been confirmed in scleractinians, thus it is not known if non-germline mutations are inherited—although this possibility has been debated vigorously^23–25^. Scleractinans are foundational species in tropical reefs and are ecologically and economically important species. Due to recent global declines, the persistence of corals in the face of climate change is uncertain. Thus, understanding whether non-germline mutations are inherited in scleractinians and how this contributes to their adaptive potential is relevant to predict their response to further climate change. Here we show that the endangered Caribbean coral *Acropora palmata* passes post-embryonic genetic mutations to its offspring, overturning the commonly-accepted view that acquired genetic variation is not heritable in animals^1^.

**Figure 1.**
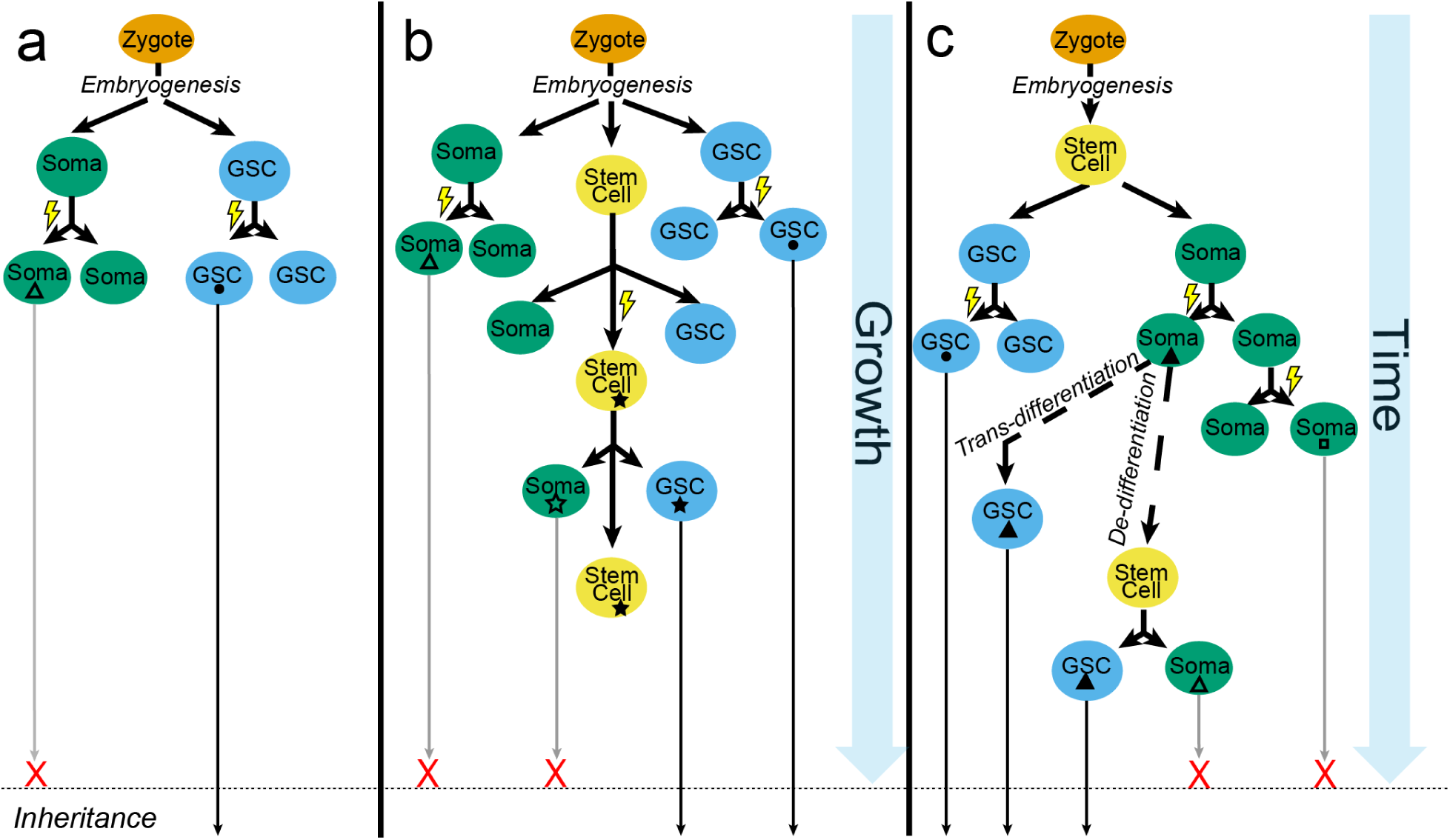
Overview of post embryonic mutations (PEMs) in animals. **(a)** Most bilaterian animals segregate germline cells from somatic cells early in development, thus preventing somatic PEMs from being inherited by offspring. **(b)** Planarian flatworms, sponges, and potentially some cnidarians continuously segregate a germline and somatic tissue from a population of stem cells as they grow, allowing for an accumulation of germline mutations that are heritable. **(c)** Occasionally, somatic cells may trans or de-differentiate into germline cells thus passing on PEMs that are somatic in origin. For all panels, colored ovals represent cell types and shapes within an oval (triangles, dots, squares) represent PEMs. Solid-fill shapes represent mutations that are inherited; outlined shapes represent mutations that are not inherited. GSC = Germline stem cell; Soma = somatic tissue. Lightning bolts are mutation causing events.

A single, sexually produced *Acropora palmata* polyp can grow into a large genet with many member colonies (ramets) via the asexual processes of polyp budding and colony fragmentation. We previously showed that *A. palmata* genets frequently harbor post-embryonic mutations that are restricted to only a subset of ramets^19^. During *A. palmata* spawning in 2017, we crossed gametes of two *A. palmata* genets from Florida, both with known post-embryonic mutations19 (Fig. 2a). Larval genotypes were analyzed at five microsatellite loci19,26 (Fig. 2a; Table S1). While most of the larvae analyzed (*n*=38, 61.3) were produced with genetic contributions from both parents, 38.7% (*n*=24) contained genetic contributions from only one parent, a surprising result because *A. palmata* has been characterized as a self-incompatible hermaphrodite^26,27^. During spawning, *A. palmata* colonies release gametes in buoyant bundles of eggs and sperm that break up at the sea-surface, and fertilization of eggs typically requires non-self sperm^28^. However, emerging evidence for deviations in coral sexual reproduction, such as parthenogenesis or self-fertilization (selfing), has been noted^26,29–32^. The exact mechanisms that lead to uniparental offspring are unknown, but this could result from the breakdown of the self-incompatibility system and/or meiotic pathways that are independent of fertilization (Fig. 2b). Among the uniparental larvae, a subset (*n*=6, 25%) inherited one known post-embryonic mutation from their parents (Table S1). At a subset of loci, most uniparental larvae (*n= 22*, 35.5%; Table S1) inherited two different alleles from one parent, indicating that the larvae were diploid. However, we designated two uniparental larvae as haploid (*n=2*, 3.2%) because they were homozygous at all loci. However, homozygous diploids and hemizygous haploids cannot be confidently distinguished from microsatellite chromatograms alone.

**Figure 2.**
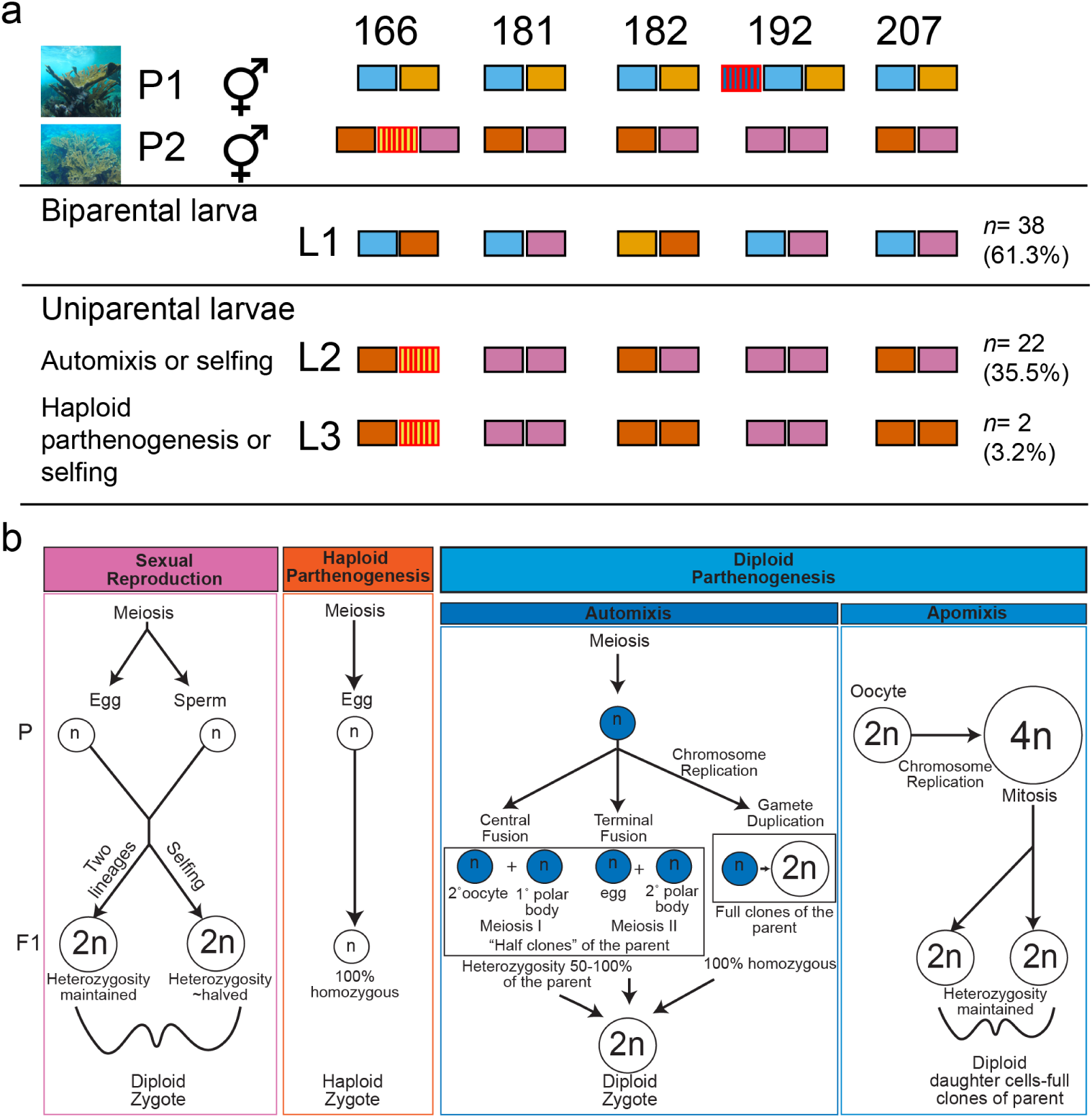
Inheritance of post-embryonic mutations. (a) Three distinct patterns of allelic inheritance were observed across five *Acropora*-specific microsatellite loci. Examples of these three patterns (L1, L2, L3) are depicted here for a subset of the samples analyzed. Gametes were collected from two hermaphroditic *Acropora palmata* colonies (P1 and P2) that each had known somatic mutations (ancestral alleles are indicated by solid colored blocks, mutated alleles are indicated by blocks with vertical lines) in one of the five loci assayed (166, 181, 182, 192, 207). While diploid in the ancestral state (represented as two blocks per locus), *A. palmata* ramets may gain alleles over time via gene duplication (represented as three blocks per locus19). P1 and P2 were crossed to produce coral larvae. Allelic patterns in most larvae (L1) followed Mendelian expectations: larvae inherited one allele from each parent at each locus (biparental larvae, 61.3%). However, allelic patterns in the remaining larvae (L2 and L3) indicated that they were uniparental in origin. L2- and L3-type larvae inherited alleles from only one parent (P2), including the somatic mutation at locus 166. L3 larvae could either be the result of haploid parthenogenesis or selfing because microsatellite analysis cannot distinguish between homo- and hemizygous states. (b) Summary of sexual reproductive strategies evident in this study and the genetic consequences of each on ploidy (n) and heterozygosity.

The following year, we observed high rates of cell division in *A. palmata* eggs collected from a single colony in Curacao, despite thorough removal of parent-colony sperm and before addition of donor sperm (Fig. 3a). Because *A. palmata* is thought to be self-incompatible, this indicated possible parthenogenesis, selfing, or sibling-chimerism at the colony level (Fig. 2b). Hundreds of these eggs developed into larvae that settled normally, took up symbionts, and matured into multi-polyp juveniles. After four months, they were preserved for single nucleotide polymorphism (SNP) analysis. For genotyping, we collected tissue samples of the parent colony (*n=10*) and one sample from each of the five nearest neighbor colonies, which could have been potential donors of contaminating sperm (Fig. 3a). DNA was extracted from thirty recruits and from all parent and surrounding colony samples (*n=15*) and submitted for SNP analysis (coralsnp.science.psu.edu/galaxy/^33^). Genotypes were assigned for 19,694 SNP probes. A multi-locus genotype analysis revealed that the five nearest neighbor colonies were ramets of the same genet as the parent colony (Table S2), with an average pairwise genetic distance of 0.0041 ± 0.0003 within a colony, and 0.0056 ± 0.0030 between colonies (Table S3; range from 0.0013 to 0.0133;^33^). All genet samples were therefore considered to be possible sources of the post-embryonic mutations found in the juveniles. The average pairwise genetic distance between genet samples and the juveniles was 0.0361 ± 0.0064 and among juveniles was 0.0508 ± 0.0082 (Table S3), similar to previous estimates for siblings^33^.

**Figure 3.**
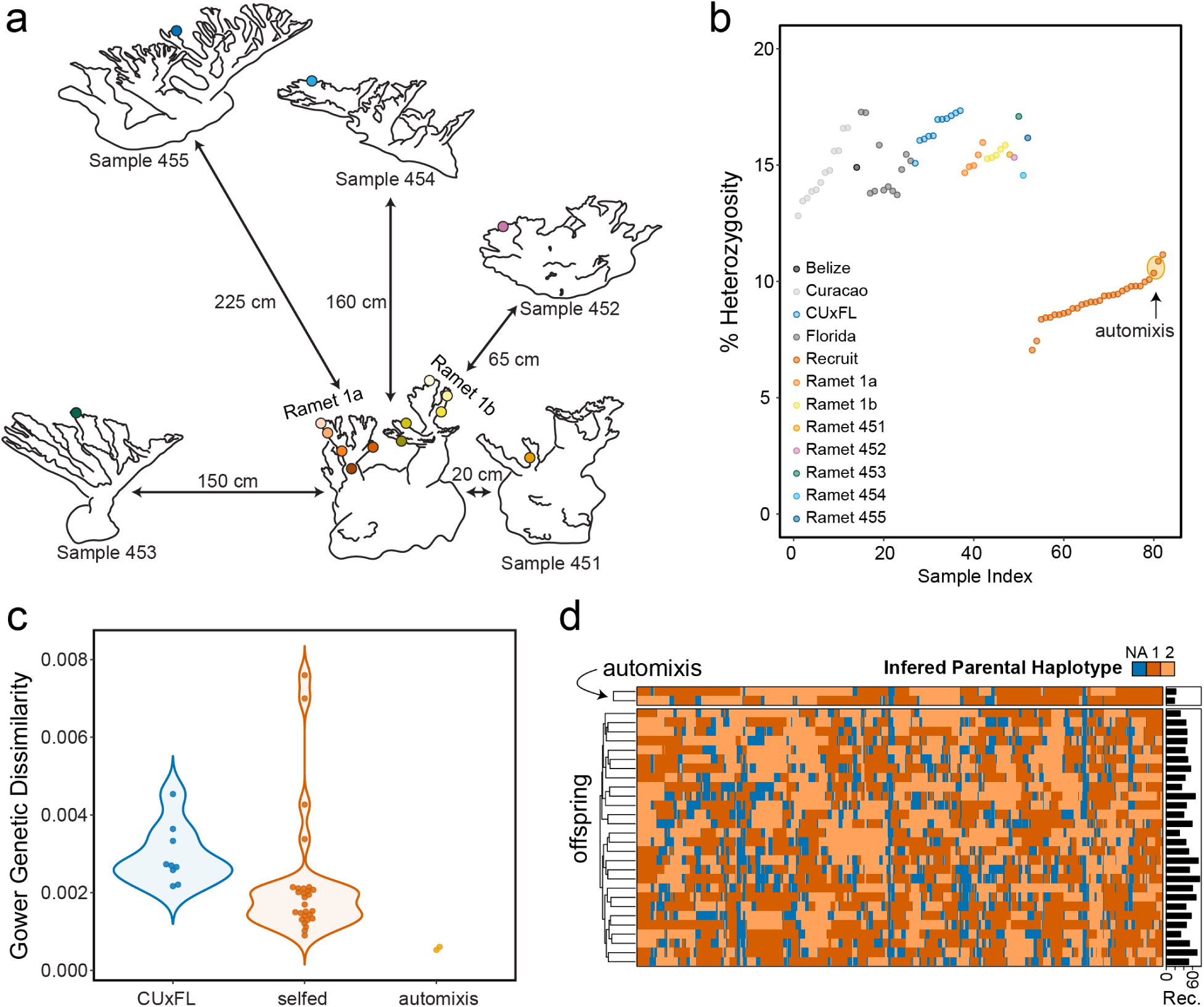
Novel sexual reproductive modes observed in *A. palmata*. (a) Sampling map of parent colony and surrounding ramets. Ramets are arranged as they were sampled on the reef and numbers next to arrows indicate the physical distance between them. Colored dots indicate the location of tissue samples analyzed via the SNP genotyping array. (b) Percent heterozygous loci (n=19,694 loci) for parent, ramets, offspring, and other Caribbean samples from Curacao, Belize, Florida and larvae resulting from a cross between Curacao and Florida colonies (CU x FL34). (c) Gower genetic dissimilarity of shared homozygous loci for all possible parental-offspring combinations of the CU x FL cross as well as selfed and automictic larvae from parent colony 441-1a. (d) Inferred haplotype blocks inherited by sibling offspring from the parent using scaffold resolution of the *A. digitifera* reference genome, revealing the number of recombination events per offspring. Phased parental haplotypes are from sister chromatids 1 (red) or 2 (orange), or could not be determined (NA, blue).

To evaluate whether juveniles were bi- or uniparental in origin, their heterozygosity was compared to adult *A. palmata* sampled from across the Caribbean, which have heterozygosity values ranging from 12.5-17.1% (Fig. 3b and Table S2;33). Curacao parent samples were within this range whereas heterozygosity in the juveniles was lower, from 7.05-11.5%. This strongly suggests that juveniles were uniparental in origin (Fig. 2b). Given that the eggs from all nights were exposed to self-sperm within the egg/sperm bundles released by the parent colony, we calculated pairwise Gower dissimilarity (GD) of shared homozygous loci for all possible parental-offspring combinations to determine if eggs were fertilized by their own sperm (selfed) or the result of parthenogenesis. GD ranges from 0 to 1, where values of 0 indicate perfect genetic identity at all loci which is expected for ‘true’ parent-offspring triads^35^. However, in practice, GD values for parent-offspring triads tend to be greater than zero due to technical errors in genotyping. We identified two recruits with mean GD of 0.0006 ± 0.0006 to putative parental donors, which is approximately a fifth of the mean GD of biparental recruits from an experimental cross between Curacao and Florida *A. palmata* gametes^34^ (Fig. 3c). These two juveniles were also assigned to the same multi-locus genet as the parent samples (Table S2), indicative of parthenogenetic origins. However, they were not full clones of the parent. The genetic distance of the two offspring that share a genet ID with the parent ramets is nearly four times that of the genetic distance between parent ramets, but a little less than half the genetic distance of the other offspring (Table S3). This may indicate that they are “half-clones” of automictic origin via central or terminal fusion after meiosis (Fig. 2b). The remaining 28 recruits had an average GD of 0.0022 ± 0.0016 similar to the biparental outcross.

We were able to assign putative parentage of the juveniles to the parental ramets (Table S4). The most common combination of parents was parents 453 and 441-1a (*n*=14 triads) followed by parents 453 and 446-1b (*n*=4 triads). The presence of several full-siblings from the same parental genet made it possible to phase the scaffolds of each sibling to further examine their reproductive mode of origin. Despite only having scaffold resolution of the *A. digitifera* reference genome, we were able to infer haplotype blocks of the siblings inherited from the parent, thereby revealing recombination events (Fig. 3d). The two parthenogenetic recruits had the lowest number of inferred recombination events (*n*=19 and 22; Fig. 3d), suggesting at least one round of meiosis occurred. The selfed recruits had on average 50.67 ± 12.20 inferred recombination events (Table S5). Although one selfed recruit had higher heterozygosity than the automictic recruits (Fig. 3b), this was not necessarily indicative that it belongs in the same category. For example, it has been shown with mathematical models that automictic individuals can share anywhere from around 50% to 100% of the heterozygous loci with the parent^36^. In addition, the automictic recruits went through less than half the average number of recombination events than the selfed recruits, consistent with both the oocyte and spermatogonium chromosomes undergoing recombination before selfing as opposed to only the oocyte chromosomes recombining before fusion of meiotic products. Combined with their low heterozygosity, these GD values and inferred recombination events suggest that there are two recruits of automictic origins and 28 recruits that are the product of selfing (Fig. 3b-e).

For genotyping probes without missing data in the parent genet (*n*=16,748), allele calls were tallied for all tissue samples from the parent genet (*n*=15) and these were compared to allele calls in the juveniles (Table S6). Because ancestral alleles should be the most common genotype among the 15 parent samples, minority SNP calls (present in 7 samples or fewer) were classified as post-embryonic mutations (*n*=271). Alternatively, minority SNP calls could also be due to technical error. We previously estimated the technical error rate of our SNP array to be 0.52% (Table S7^33^) and the number of detected mutations (*n*=271) was three times higher than the expected number of technical errors (*n*=104). The number of inherited mutations in juveniles (*n*=139) also exceeded the error rate and we further validated SNP calls via RFLP (see below). Most mutations (72.3%) were due to transitions and fewer (27.7%) were due to transversions (Table S8). Putative mutations were found in both protein-coding (13.28%) and non-protein-coding regions (86.72%, Table S9). Parent 453 had the most mutations (*n*=149, Table S2) followed by parent 448 (*n*=52), a part of ramet 1b from the parent colony (Fig. 3a). Out of these mutations, 139 were detected in at least one juvenile and the number of mutations per juvenile ranged from 11 to 50 (Fig. 4a, Table S2 and S6). Loci with putative inherited post-embryonic mutations most often changed from a homozygous state to a heterozygous state (gain of heterozygosity (GOH), *n*=91), followed by loci that went from a heterozygous state to a homozygous state (loss of heterozygosity (LOH), *n*=45, Table S8). While all but one LOH mutation was shared between a parent and offspring, less than half the GOH mutations were shared. Elevated GOH identified in the parental samples could be due to sample quality, which has been shown to inflate heterozygosity calls with genotyping array data^33^.

**Figure 4.**
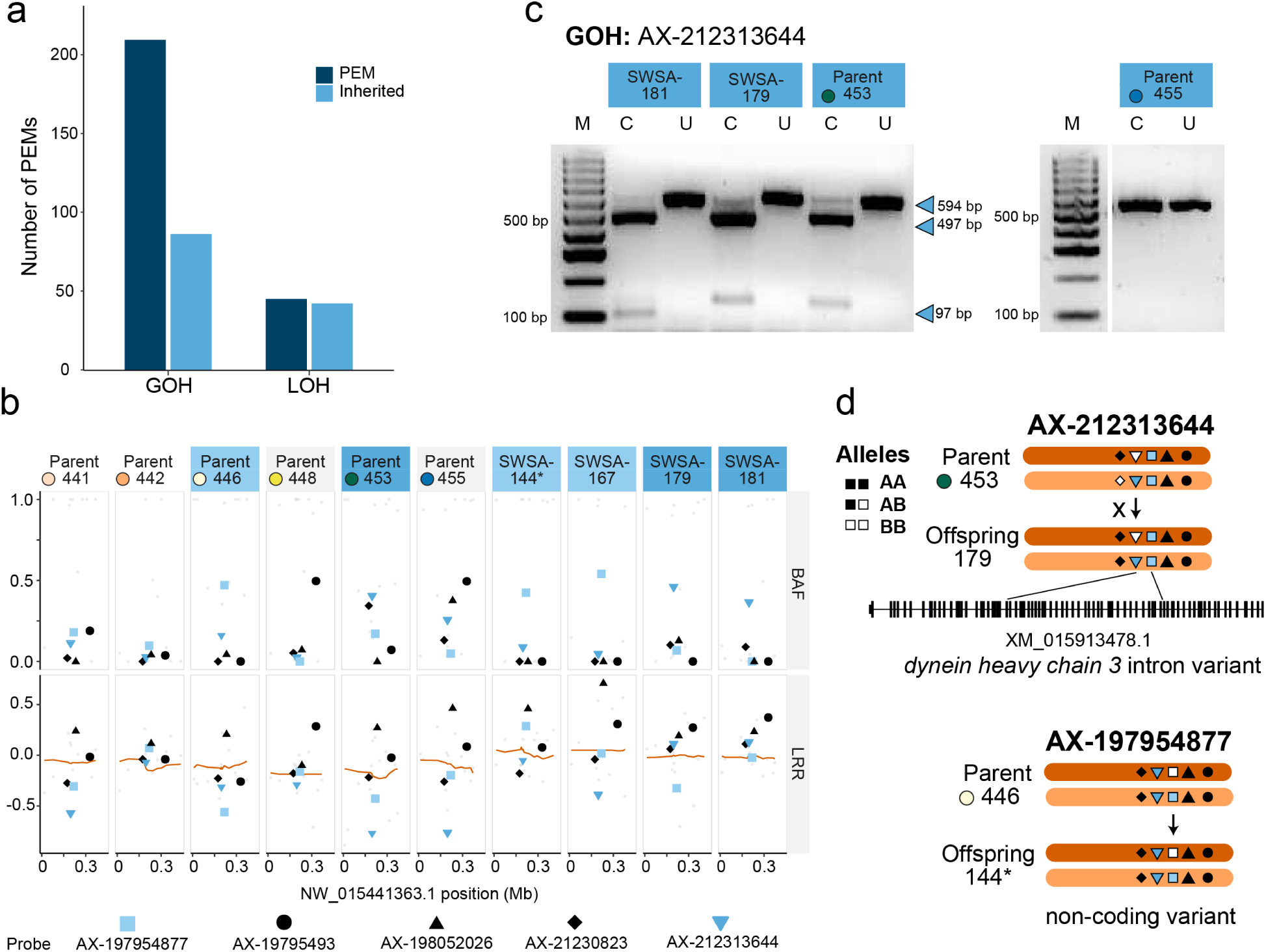
Characterization of inherited post-embryonic mutations (PEMs) detected in the SNP data. **(a)** Gain of heterozygosity (GOH) mutations outnumbered loss of heterozygosity (LOH) mutations. Proportionally, more LOH then GOH mutations were inherited. **(b)** Maps of two metrics of allele signal intensity, the BAF (total allelic intensities) and LRR (relative allelic intensities) that are commonly used to investigate somatic mutation and copy number variation; orange line represents sliding average of 20 SNPs along the scaffold (position in megabases Mb, x-axis). **(c)** RFLP validation of inherited GOH mutation, AX-212313644. For each sample, the uncut (U) and cut (C) PCR products are shown. Each gel contains a size standard (lane M, bp = base pairs). The two recruits (SWSA-179 and SWSA-181) share the heterozygous mutation with the parent 453, resulting in 3 bands, while the parent 455 predicted to have the non-mutant homozygous state for this site did not cut, resulting in 1 band **(d)** RFLP-validated post-embryonic mutation, AX-212313644 is found within the intron of the dynein heavy chain. GOH mutation, AX-197954877 is found on the same scaffold as AX-212313644. Symbols represent the five mutations detected along this scaffold in at least one sample. Black shapes: not inherited, blue shapes: inherited, filled shapes: A allele, and open shapes: B allele. Arrows denote inheritance and crosses do not. One automictic offspring is denoted by an asterisk.

Mapping two metrics of allele signal intensity, the b-allele frequency (total allelic intensities) and log R ratio (relative allelic intensities) that are commonly used to investigate somatic mutation and copy number variation, confirmed that most mutations in the diploid juveniles were not a product of copy number differences (Fig. 4b).

To validate the detected SNP mutations, SNP-RFLP markers were designed for 19 of the inherited loci^33^ (Table S10). SNP-containing regions were amplified by PCR, and the resulting PCR product was digested by a restriction enzyme (Fig. 4c). Markers that produced both sharp PCR bands and clear results in restriction digests were further investigated. Of these markers, GOH variant mutation locus AX-212313644 produced the clearest banding patterns on gels. Two larvae (SWSA-179, SWSA-181) share the heterozygous mutation with the parent 453, resulting in 3 bands (Fig. 4c) while parent 455 predicted to have the non-mutant homozygous state for this site did not cut, resulting in 1 band (4c). The RFLP marker confirms the inheritance of the GOH mutation found at this locus discovered in the SNP array data. Post-embryonic mutation locus AX-212313644 is a non-coding variant found within the intron of the *dynein heavy chain 3*, upstream of another intron mutation in the same parent sample but not shared with the other juveniles screened in this study (Fig. 4d).

By investigating coral juveniles from parents with known post-embryonic mutations, we provide evidence for the inheritance of post-embryonic mutations (possibly somatic in origin) in animals. Because coral genets can persist for hundreds to thousands of years, post-embryonic mutations can rise to high frequency in parts of a genet as a result of selection and/or stochastic processes^37^. These mutations could then be dispersed over shorter distances by fragmentation, and over longer distances by pelagic larvae that have inherited the mutations.

The origin of post-embryonic mutations in the adult *A. palmata* remains unknown. Mutations identified here as being passed from an adult colony to meiotically-produced offspring (Fig. 2 and 4) may have originated in stem cells, which were passed on to the germline during differentiation (Fig. 1b). The mutations may have also originated in the soma, de-differentiated into stem cells, and then re-differentiated into germ cells. Alternatively, if scleractinian corals develop like most bilaterian animals (Fig. 1a) and segregate a germline early in development, most of these mutations may have originated in the somatic tissue and transdifferentiated from soma to germ cells (Fig. 1c). In either case, the adult mutations must have occurred post-embryogenesis, because they were not shared among all ramets of the parent genet. Buss ^13^ posited that somatic post-embryonic mutations, if heritable, might be beneficial to a modular organism. Immediately after a somatic cell mutates, it undergoes somatic environmental selection and its propagation depends on successfully outcompeting other somatic lineages for positions in the germline^38^. Beneficial (or neutral) mutations that survive post-embryonic environmental selection may thus be disproportionately represented in the cells of a genet^39^, an advantage that ancestral germ line mutations do not have. Studies of *Acropora cervicornis* growth show that new polyp tissue (including gonads) appears to be differentiating from epidermal or somatic tissue. Ten of the uniparental offspring carried the same mutation (SNP AX-212312351) as a single parent sample (453), indicating that it may have been somatic in origin.

Curiously, the coral juveniles analyzed here had a substantially higher percentage of uniparental larvae (msat: 38.71%, SNP: 100%) than those in previous studies (ca. 1-2%; ^26^,^27^). In the Caribbean, failure of sexual recruitment is prevalent among most important reef-building species^23^,^24^, therefore asexual processes (fragmentation and reattachment) have dominated local population growth on many reefs^40^. The majority of large, long-lived Caribbean reef-building coral species are self-incompatible hermaphrodites which require the presence of gametes from another genet for successful fertilization^27^,^41^. Thus, the physical distance between genets and/or asynchronous spawning of neighboring genets may be insurmountable obstacles for sperm to successfully fertilize spawned eggs^25^,^42^. Further, in the absence of dense populations of potential mates (and their water-dispersed reproductive pheromones), corals may prioritize the generation of uniparental larvae, as outcrossing and selfing may result in the recombination of beneficial post-embryonic mutations into different genetic backgrounds. An alternative to this coral choice hypothesis is the coral senescence hypothesis. Aging genets may lose the ability to recognize and reject self-gametes due to mutations in *e.g.*, gamete recognition or self-incompatibility systems. A third explanation for the high instance of uniparental larvae observed in this study is the environmental driver hypothesis: toxins such as endocrine disruptors are now present in reef environments^43^ and may change sexual reproductive behavior. As coral populations decline, uniparental reproduction may increase in frequency, potentially resulting in decreased genetic diversity. At the same time, there is the intriguing possibility that post-embryonic mutations might help buffer such losses of genetic diversity.

The inheritance of acquired genetic variation fundamentally changes how we understand animal adaptation. By acquiring mutations throughout their decades-to millennia-long lifespans and passing these mutations on to their meiotic offspring, corals may be able to confer and disperse favorable phenotypes. Perhaps the inheritance of acquired mutations that are beneficial and their subsequent dispersal via the mobile life stage and/or the incorporation of uniparental sexual strategies in the absence of reproductively active (or any) mates helps to explain why corals have been able to persist for over 500 million years^44^. In some *Drosophila* ^45^ and vertebrate^46^ species that are facultatively parthenogenetic, there are no clear subpopulations between individuals who reproduce automictically and sexually unless rare automixis facilitates the colonization of a sparse or unpopulated area^36^. We can only speculate on what this reproductive flexibility afforded coral populations over geologic time because reconstructing past evolutionary events is difficult. Elucidating the effect of post-embryonic mutations on gene expression and epigenetic patterning will be critical for modeling the ecological impacts of simultaneous use of different sexual strategies. Modeling the evolutionary genetic consequences of post-embryonic mutations may also help to explain their roles in mutational meltdown under Muller’s ratchet^47^ or in adaptation in the next century. Moving forward, the question remains how, where, and under what circumstances post-embryonic mutations will positively or negatively affect coral populations today.

## Methods

### 2017 mutant gamete collection and crosses

We targeted *A. palmata* colonies with post-embryonic mutations at their microsatellite loci for gamete collection. Gamete bundles were collected on August 11^th^, 2017 from two reefs in the Florida Keys: Sand Island Reef (SIR, Lat. 25.0179; Long. −80.368617) and Elbow Reef (ELR, Lat. 25.15185; Long. 80.2497). At SIR, gametes from two colonies were collected: Sand Island Blue Mutant and Sand Island Orange. At ELR, gametes were collected from the colony named Elbow Green Mutant. The first cross was between Sand Island Blue Mutant and Sand Island Orange (hereafter referred to as cross 1MP (One Mutant Parent)). The second cross was between Sand Island Blue Mutant and Elbow Green Mutant (hereafter referred to as cross 2MP (Two Mutant Patents)). For cross 1MP, we combined 1 ml of eggs and 1 ml of sperm (concentration ~10^6^) from each parent at 23:15 and waited 1.5 hours to allow fertilization to take place. Following fertilization, embryos were washed two times with filtered seawater (FSW) and placed in a 1 L container with FSW to grow overnight. For each parent, eggs and sperm from the same colony were also combined in selfing controls. After 1.5 hours, it was noted that Sand Island Blue Mutant and Sand Island Orange did not undergo any self-fertilization because no dividing embryos were observed. For cross 2MP, 1 ml of eggs and 1 ml of sperm from each mutant colony were combined at 00:38 and allowed 1 hour to fertilize. At approximately 01:38, embryos were washed two times with FSW and placed in a 1 L container with FSW to develop overnight. We performed selfing crosses for 2MP as well and noted that Sand Island Blue mutant eggs and sperm did not self while Elbow Green Mutant eggs and sperm did self, with a self-fertilization rate ranging from 0.0034 to 0.070% over two nights. The following morning, we removed any eggs that had not fertilized and performed a water change. We continued to do maintenance water changes on the crosses twice a day for three days. Generally, larvae began swimming at the end of the third day. Finally, at 96 hour post-fertilization, individual swimming larvae from each cross (1MP *n=300*; 2MP *n=200*) were preserved individually in 96% non-denatured ethanol and stored at −20°C until shipment to the Pennsylvania State University.

### Larval DNA extraction and microsatellite analysis

DNA was extracted from ninety-five larvae from each cross. The larvae were rinsed once with fresh ethanol to remove debris and ethanol was then replaced with 20 μl of 5% Chelex solution and 2 μl of 20 mg/ml Proteinase K. Samples were vortexed for 2 seconds, digested overnight at 55°C, and heated to 95°C before cooling to 4°C.

The first multiplex PCR amplified microsatellites 166, 192, and 181^26^. Microsatellite markers were amplified by PCR in a 10 μl reaction volume containing water, 10X Original Buffer, 25 mM MgCl, 10 mM dNTP, 5 μM 166-pet, 5 μM 192-6fam, 5 μM 181-ned, 5 U/μl GoTaq Flexi DNA Polymerase (Promega, WI, USA), and 1 μl of DNA template. The second multiplex PCR amplified microsatellites 182 and 207^26^. DNA was amplified in a 10 μl reaction volume containing water, 10X Original Buffer, 25 mM MgCl, 10 mM dNTP, 5 μM 182-6fam, 5 μM 207-pet, 5 U/μl GoTaq Flexi DNA Polymerase, and 1μl of DNA. Each PCR mixture was denatured at 94°C for 5 minutes followed by 35 cycles of: 94°C for 20 seconds, 54°C for 20 seconds, and 72°C for 30 seconds. Samples were held at 72°C for 30 minutes for final extension. Samples were checked for sequence amplification success by running 4 μl of PCR product on a 2% agarose gel. Once amplification was verified, samples were sent to the Genomics Core Facility at the Pennsylvania State University for Fragment Analysis on the Applied Biosystems 3730XL DNA analyzer. Two persons independently called allele sizes for each microsatellite locus using Genemapper 5.0 software (Thermo Fisher Scientific). In *A. palmata*, post-embryonic mutations manifest as additional alleles in microsatellite chromatographs. Most frequently, a mutant genotype shows one additional allele at one of the five loci. The mutation is most often one repeat size larger or smaller than the ancestral allele of that genet^19^. The ancestral allele is established by comparing at least five samples from a genet and determining the majority alleles. Through this analysis, Sand Island Blue Mutant had a third allele at microsatellite locus 166 that the ancestral Sand Island Blue genet did not have^19^. The Elbow Green Mutant genet had a third allele at microsatellite locus 192 that the ancestral Elbow Green genet did not. Post-embryonic mutations were not detected in Sand Island Orange at any of the five microsatellite loci.

### 2018 spawning collection

Seven nights after the full moon (AFM, September 2^nd^, 2018), a subset of *A. palmata* colonies from Spanish Water reef (Lat. 12.0636, Long. −68.8532) in Curacao produced eggs that underwent cell division in the presence of only their own sperm. Despite efforts to quickly separate sperm from eggs after subsequent spawning events, eggs from these colonies continued to display apparent self-fertilization, followed by normal larval development and normal larval swimming behaviors, on two additional spawning nights. Embryos from all three spawning nights were allowed to develop in containers of filtered seawater (spun polypropylene filters, 0.5 μm). Larvae were then shipped to Mote Marine Laboratory for settlement and rearing. At month four post-fertilization, 81offspring were preserved in 96% ethanol and shipped to the Pennsylvania State University for genetic marker analysis. In addition, the coral colony that produced the apparently self-fertilized eggs was sampled in five locations along two of its branches (*n=* 10). In addition, one sample was taken from each of the five nearest neighboring colonies (*n=5*, Fig. 3a). These fifteen samples were also preserved in 96% ethanol for genetic marker analysis. To collect tissue from the adult colonies, whole polyps including skeletal material were sampled. These adult colonies had spawned gametes the previous night and thus tissue samples most likely contained a mixture of reproductive and vegetative tissue.

### SNP analysis and d*e novo* detection of post-embryonic mutations

We extracted genomic DNA from 81 recruits and 15 parent samples using the DNeasy kit (Qiagen, USA) following the manufacturer’s protocol with slight modifications optimized for corals (https://doi.org/10.17504/protocols.io.bgjqjumw). Samples were genotyped using an Affymetrix genotyping array^33^ and analyzed using the STAG analysis portal (https://coralsnp.science.psu.edu/galaxy/^33^). A total of 19,694 genotyping probes were extracted for downstream analyses using vcfR package in R^48^. Genotype calls were converted into ‘0/0’, ‘0/1’, or ‘1/1’. A call of ‘0/0’ is homozygous for allele A at that locus, likewise, a call of ‘1/1’ is homozygous for allele B at that locus and a call of ‘0/1’ means that the sample is heterozygous at a locus with respect to allele A and B. Missing data and heterozygosity for each sample was calculated as the sum of probes with ‘NA” or ‘0/1’ calls, respectively, divided by the total number of probes in R^49^. Gower’s genetic dissimilarity and parentage assignment for the respective crosses were calculated using the R script *apparent* as previously described^35^. To infer parental haplotype blocks and recombination events, the genotype data from the offspring siblings were phased using the *bmh* and *recombination* functions in the hsphase R package^50^ and plotted with the ComplexHeatmap package^51^. Post-embryonic mutations in the parent samples and ramet samples (representative of possible variation within the parent colony) were tallied for each locus across parent colony and ramet samples. Based on the premise that the ancestral allele should be more frequent than the mutated or alternate allele, alleles were designated as ancestral if eight or more ramets shared the allele. Parental mutant alleles were similarly tallied in the offspring. The b-allele frequency and log R ratios were calculated using the ‘affy2vcf’ bcftools plugin (https://github.com/freeseek/gtc2vcf), as part of the STAG workflow. Genomic location and predicted effect of the mutations were found with snpEff v4.3^52^ using the *A. digitifera* genome assembly^53^.

### RFLP validation of post-embryonic mutations

SNP-RFLP markers were designed using methods described in Kitchen, et al. ^54^ with modifications outlined below. Briefly, 28 of the 139 mutations were screened using WatCut (http://watcut.uwaterloo.ca/template.php?act=snp_new). Primers were designed using Primer 3^55^ based on 500 bp of flanking sequence around the SNP extracted with bedtools *getfasta* utility v2.27.1^56^ from the *A. digitifera* genome^53^ for 19 of the 28 variant mutations with restriction enzyme cut-sites identified (Table S10). DNA of two of the larvae (SWSA-179, SWSA-181) and two parents (453 and 455) were chosen for validation of their SNP calls. SNP-containing regions were amplified by PCR in a 10 μl reaction volume containing water, 1X NH4 Buffer (Bioline, Boston, MA), 3 mM MgCl (Bioline, Boston, MA), 1 mM dNTP (Bioline, Boston, MA), 250 nmol forward and reverse primers (IDT, Coralville, Iowa), 1 unit Biolase Taq (Bioline, Boston, MA), and 1 μl of template DNA. PCR product was then denatured at 94°C for 5 minutes, then 35 cycles of PCR were performed (94°C for 20 seconds, 55.2°C for 20 seconds, 72°C for 30 seconds). PCR wells were then held for a final extension at 72°C for 30 minutes. The resulting PCR product was then directly digested by restriction enzyme HypAV in a 10 μl reaction volume containing 5 μl PCR product, 1X CutSmart Buffer, 0.5 μl of enzyme HpyAV (New England Biolabs, Ipswich, MA), and water to volume. Digests were held at 37°C for 1 hour, then held at 65°C for 20 minutes to heat-kill the enzyme. PCR and digest products were visualized using electrophoresis on 2% agarose gel.

## Acknowledgements

We would like to thank Kathryn Stankiewicz for her assistance with R and Drs. Dana Williams and Margaret Miller provided field support for the 2017 experiment. We also thank Meghann Devlin-Durante for her help with the microsatellite analysis, Dr. Mary Hagedorn for overseeing the larval shipment to Mote Marine Laboratory, Daisy Flores, Lucas Tichy, and Valerie Chamberland for dive and lab assistance, and Cornelia Osborne for manuscript edits. Funding was provided by Funding for this project was supported by NOAA Office for Coastal Management NA17NOS4820083, and NSF OCE-1537959 to IBB. Special thanks to the Paul G. Allen Family Foundation for funding spawning operations and larval rearing in Curacao and Florida 2018.

## Data and code availability

SNP data for the samples in this study can be exported at https://coralsnp.science.psu.edu/galaxy/. Code will be provided on a GitHub repository at the time of publication.

## Author contributions

K.L.V.K. carried out the 2017 experiment, designed the 2018 experiment, analyzed data, and co-wrote the paper; S.A.K. contributed to the design and execution of the 2017 experiment, carried out part of the SNP analyses and co-wrote the paper; T.L.C. assisted with the parentage analysis and performed the RFLP assays; S.A.V. helped with the SNP analyses and edited the paper; A.N.C. assisted with the 2017 experiment and edited the paper; C.P. reared the recruits used for SNP analyses; K.L.M. and M.J.V. collected spawn and tissue samples, coordinated fieldwork, and edited the paper; I.B.B. oversaw the research, co-designed the 2017 and 2018 experiments, obtained funding, and co-wrote the paper.

## Additional Information

**Supplementary information** is available for this paper. Tables S1 through S10 are compiled into an excel document.

